# Scalable and cost-efficient custom gene library assembly from oligopools

**DOI:** 10.1101/2025.03.22.644747

**Authors:** Chase R. Freschlin, Kevin K. Yang, Philip A. Romero

## Abstract

Advances in metagenomics, deep learning, and generative protein design have enabled broad in silico exploration of sequence space, but experimental characterization is still constrained by the cost and scalability of DNA synthesis. Here, we present OMEGA (Oligo-based Multiplexed Efficient Gene Assembly), a low-cost, accessible method for assembling hundreds to thousands of full-length genes in parallel using standard laboratory techniques. OMEGA computationally fragments target genes into short, high-fidelity Golden Gate-compatible oligonucleotides that can be ordered as a pooled library and assembled across multiplexed subpools. We systematically optimized the number of fragments per gene and orthogonal ligation sites per reaction and determine that OMEGA can assemble up to 2.6 kb constructs using as many as 70 Golden Gate sites. To validate the approach, we assembled and functionally screened a library of 810 natural and synthetic GFP variants, recovering 94–97% of target sequences with high uniformity. OMEGA enables precision library construction at scale, with per-gene costs as low as $1.50, and offers a broadly applicable solution for bridging computational protein design with high-throughput experimental validation. We have developed OMEGA as an open-source software package and an easy-to-use Colab notebook available at https://github.com/RomeroLab/omega.

## Introduction

The explosion of biological data, coupled with advances in deep learning and generative AI, has transformed protein engineering into a data-driven discipline. We have unlocked vast sequence diversity with advancements in large-scale metagenomic sequencing^1,2^ and generated detailed sequence-function landscapes with high-throughput data generation.^3–5^ These data have given us the power to design limitless mutants, synthetic homologs, and de novo designs in silico that explore vast swathes of sequence space.^6–9^ However, the ability to experimentally characterize these designs remains a major bottleneck. Scalable gene synthesis methods are essential to bridge this gap, enabling targeted exploration of sequence space with the same precision as data-driven models to expand our knowledge of protein function.

Current DNA synthesis methods are constrained by a strict trade-off between sequence length and scale that limit the scope of precision libraries. To synthesize custom sequences, researchers must choose between synthesizing a few long sequences as gene fragments (<=5000 bp) or many short sequences as oligopools (∼300 bp) from manufacturers like Twist Bioscience and IDT. Oligopools are heterogenous mixes of single stranded DNA where each strand is uniquely specified by the user.^10^ They are significantly cheaper than fragment synthesis per base, but constrained to DNA fragments shorter than most protein coding genes.

We can expand the encodable length of oligopools by using DNA assembly to build larger constructs from multiple oligos. Scaling DNA assembly for large, diverse sequence libraries presents challenges in workflow complexity, specialized equipment needs, assembly accuracy, and construct length. For example, Polymerase Cycling Assembly (PCA) can stitch together multiple oligos in individual reactions but struggles when scaled to many highly similar sequences.^11–15^ High-throughput PCA assemblies using DropSynth^16^ resulted in low-fidelity libraries that contained ∼80% of target genetic constructs with a high background of incorrect products. Alternatively, Golden Gate cloning uses Type IIS restriction enzymes to generate unique DNA overhangs called Golden Gate (GG) sites (commonly 4 bp), which allows for efficient assembly of multiple DNA fragments.^17^ Data-optimized Assembly Design (DAD) identifies combinations of GG sites that yield high-fidelity DNA assemblies, allowing for high-complexity assemblies using tens of GG sites, and have enabled the assembly of large constructs directly from oligopools.^18–20^ In theory, this approach can be scaled to recover multiple constructs from a single Golden Gate assembly if all GG sites are sufficiently orthogonal between target sequences.

In this work, we present a low-cost and accessible method for precisely assembling hundreds to thousands of genes in parallel using standard molecular biology techniques and equipment. Our method computationally splits a list of target genes into fragments optimized for high-fidelity GG assembly and that can be ordered as an oligopool. The oligopool is divided into 10-100 subpools in a microtiter plate, where several genes are assembled per pool, yielding hundreds of precisely specified genes. To develop this method, we systematically optimized key DNA assembly parameters, including the number of GG sites per subassembly and the number of ligated fragments, to assess scalability across gene lengths and library sizes. We demonstrated the scalability of our approach by assembling a diverse panel of 810 natural and designed Green Fluorescent Protein (GFP) genes, achieving 94-97% recovery of target genes. The final gene library was highly uniform with 87-92% of genes within tenfold of the median-abundant sequence. We performed downstream functional screening of the designed GFP variants to demonstrate the quality of the assembled genes. Our method requires only standard lab equipment, minimal hands-on time, and takes two days start-to-finish, enabling the cost-effective assembly of hundreds to thousands of genes for $1.50 to $14 per gene for constructs up to 2.6 kb.

## Results

### OMEGA: Oligo-based Multiplexed Efficient Gene Assembly

Most proteins–including many with applications in medicine and chemistry–require individual synthesis as gene fragments because their genes are too long to fit on a single oligonucleotide (**Fig. 1a**). Proteins such as PETases, luciferases, and Cas9, which can be engineered to recycle plastics, image in vivo processes, and edit genomes, cost more than $50 USD per variant to synthesize, making large-scale experiments impractical. There is significant interest and need for new methods to assemble DNA fragments from oligonucleotide pools (oligopools) into thousands of gene-length sequences.^21^

**Figure 1.**
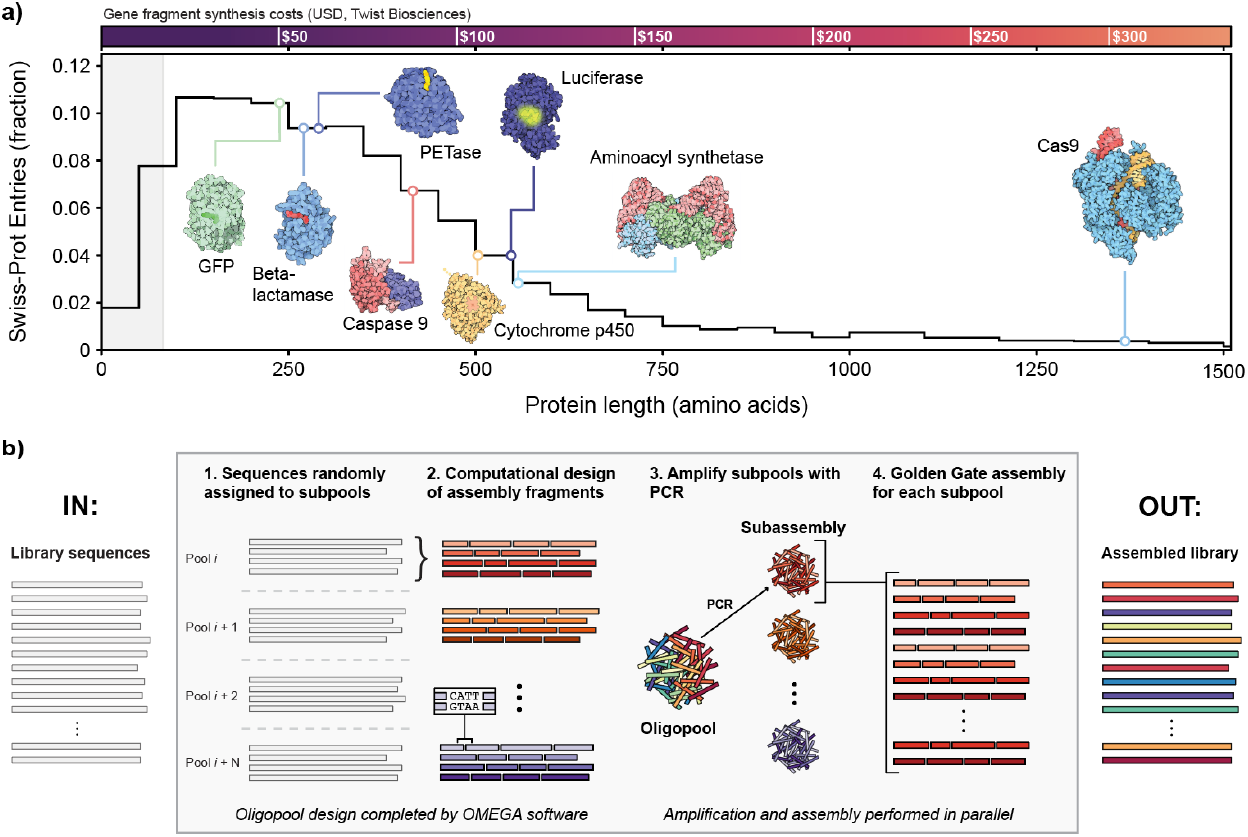
Oligo-based Multiplexed Efficient Gene Assembly (OMEGA). **(a)** Distribution of protein lengths for all entries in Swiss-Prot. Most proteins are too large to encode on single oligos, represented as the shaded grey area. Oligos provide approximately 250 bp of coding sequence given a 300 bp oligo with PCR and cloning adaptors. The synthesis cost per gene is shown above the distribution using Twist Bioscience prices for fragment synthesis. Structures are from RCSB Molecule of the Month articles and use the following structure codes from left to right: 1GFL, 4EYL, 5XH3, 1NW9, 1W0E, 2D1S, 1ASZ, 4OO8. **(b)** OMEGA design and assembly workflow. Users provide a list of codon-optimized sequences that OMEGA randomly sorts into N subassembly pools. For each subpool, OMEGA designs fragments that use high-fidelity GG sites and ensure that all pieces are within oligo size constraints. Subassembly pools are separately amplified and assembled into full-length gene products in parallel, after which assembly products are combined to recover the designed library.

We aimed to develop a simple and accessible method for assembling 100s-1000s of genes from oligopools that reduces synthesis costs and scales with library size. To assemble genes from oligopools, a full-length gene must be broken into fragments that 1) are small enough to encode on an oligo and 2) possess a unique GG site at the 3’ and 5’ ends. There are many fragmentation patterns that satisfy these constraints, but not all of them use GG site combinations predicted to assemble with high-fidelity. We use DAD to formulate gene fragmentation as an optimization problem, aiming to identify fragmentation sites that maximize GG assembly fidelity in a multiplexed reaction, where multiple genes are assembled simultaneously using unique orthogonal overhangs. The result is a pooled assembly that recovers individual genes due to the high-fidelity GG sites uniquely assigned to each gene. We call this approach OMEGA for Oligo-based Multiplexed Efficient Gene Assembly. OMEGA is inspired by recent work by Pryor et al.^18^ demonstrating the capacity of GG cloning to achieve complex DNA assemblies with up to 52 DNA fragments with DAD.

We present OMEGA as a software package (https://github.com/RomeroLab/omega) that optimizes gene fragmentation for high-fidelity Golden Gate assembly. It generates a list of fragments orderable as an oligopool (**Fig. 1b**) that can be efficiently assembled using standard lab equipment and established GG cloning protocols. Users input a set of codon-optimized target genes, which can be both sequence-diverse and variable in length. The library of *n* genes is randomly sorted into *m* subpools and OMEGA designs gene fragments for each subpool. Each subpool is indexed by unique primer pairs that selectively amplifies each subpool for GG assembly in parallel on a microtiter plate.^22^ The size of each subpool is determined by the total number of GG sites used per reaction (set by the user) and gene length. Finally, the subpools are combined to create a comprehensive gene library ready for downstream functional screening and characterization.

### Probing the limits of GoldenGate multiplexing for scalable gene assembly

OMEGA’s scalability depends on the degree of multiplexing within each subpool. Higher multiplexing allows more genes to be assembled per subpool and supports the assembly of longer genes composed of additional oligo fragments. The upper limits of multiplexing are determined by the number of unique GG sites that can be used without significantly reducing assembly fidelity. As the number of GG sites in a reaction increases, the risk of misligation between non-matching sites rises, leading to a decline in assembly accuracy. The maximum number of GG sites that can be included in a single reaction while maintaining high efficiency and fidelity remains largely unknown, in part because GG assemblies are done under ideal conditions not reflected here. We designed several oligopool assemblies that mimic OMEGA conditions to systematically explore how set size and gene length affect both the size of an OMEGA library and the length of genes we can assemble.

To assess the impact of multiplexing on assembly fidelity, we used OMEGA to design assemblies with varying numbers of GG sites per reaction. We performed two-fragment assemblies, scaling the number of GG sites from 20 to 70 by adjusting the number of two-fragment constructs per reaction. For example, we assembled 40 fragments to test a set of 20 GG sites and 140 fragments to test 70 sites. In addition to the designed GG ends, each fragment included a unique 25mer sequence, allowing us evaluate assembly fidelity using Illumina sequencing. For each set size, we performed two assemblies using diverse sets of GG sites to evaluate variability between solutions (**Supplementary Fig. 1**). We then performed the OMEGA protocol by separately amplifying each subpool, performing GG assembly with a high-fidelity protocol. We analyzed fidelity, coverage, and bias using deep Illumina (**Fig. 2a**) for all assemblies.

**Figure 2.**
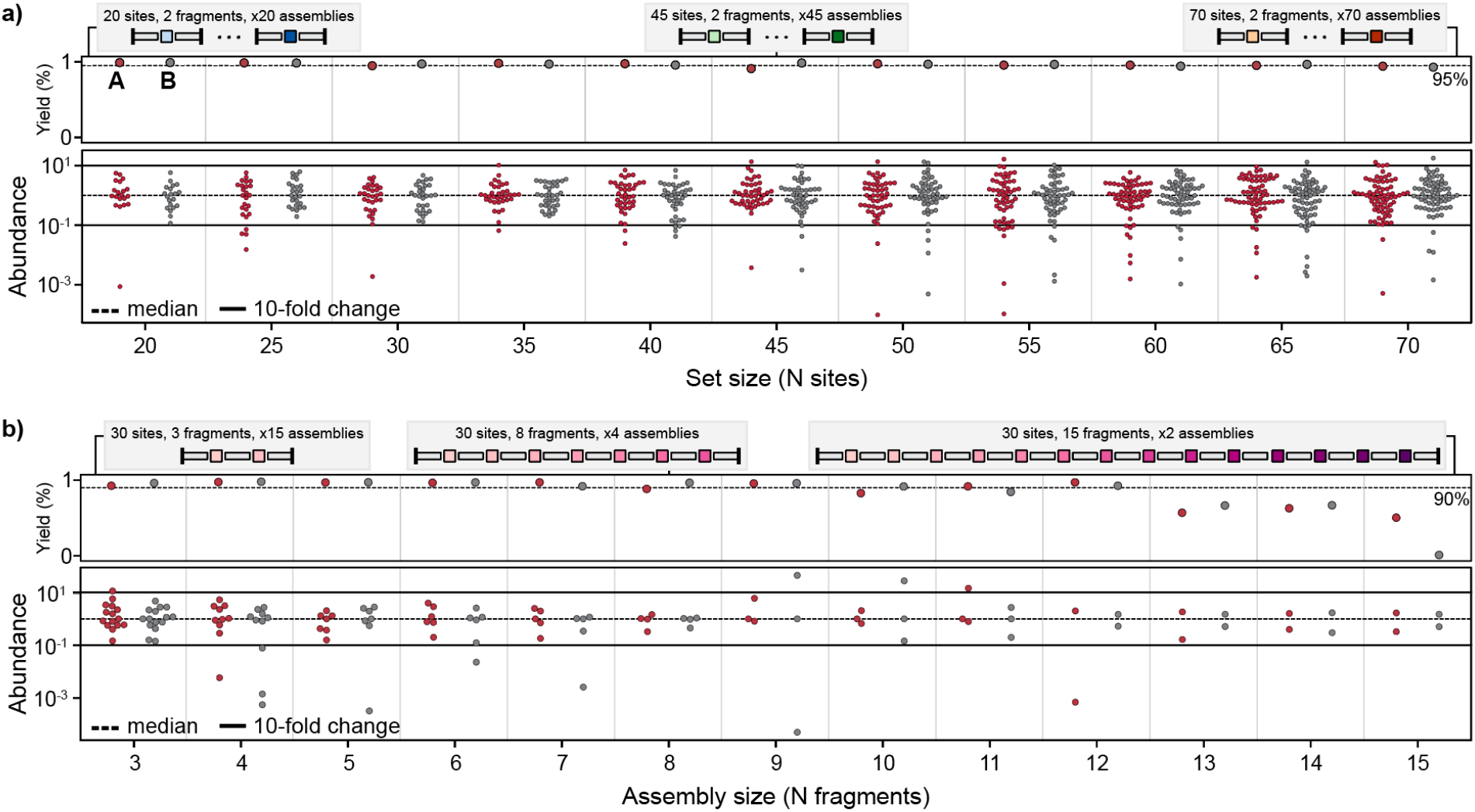
Fidelity of Golden Gate assemblies of varying complexities and sequence lengths. Target sequences consist of 25mer fragments connected by a GG site assembled into a destination cassette. All trials were assembled using a high-fidelity Golden Gate protocol (details in methods) and sequenced with more than 26,000x coverage. Yield represents the number of NGS reads that are correct assemblies versus all reads. Abundance is represented as x-fold from the median abundant sequence. We consider sequences to be over- or under-represented if they are more than 10-fold from the median. **(a)** Fidelity of assemblies using 20 to 70 GG sites in increments of 5. Red and grey dots represent two independent design trials. Most assembly sizes achieve >95% yield with a handful of under-represented sequences. Sequences are rarely over-represented. **(b)** Fidelity of assemblies with sequences using 3 to 15 fragments. Product yield remains constant around 90% until 13 fragments, at which point yield drops to ∼65%. The distribution of sequences are typically within 10-fold of the median.

Across all tested GG set sizes (20 to 70 sites), over 90% of sequenced assemblies were correct with most assemblies exhibiting >95% correct product. With the exception of two subpools, all target sequences were observed at least once. One 40 site and one 70 site subpool assembled 97% and 98% of target sequences. We assessed correct assemblies based on fragment order and did not consider sequence-correctness. Thus, these errors reflect the impact of misligation events on assembly fidelity. The assembly products were highly uniform across set sizes, with 92.6% falling within tenfold of the median-abundant sequence on average. While underrepresented sequences became more common as set size increased, overrepresented sequences remained rare across all conditions. Construct abundance was not significantly affected by GG site ligation efficiency or the abundance of individual fragments after PCR (**Supplementary Fig. 2 and Supplementary Fig. 3**).

Next, we investigated how construct length affects OMEGA’s assembly fidelity by determining the maximum number of fragments that can be reliably assembled. Longer genes composed of more fragments will decrease assembly fidelity through increased partially assembled products due to the cumulative effect of multiple imperfect ligation events. We used OMEGA to design assemblies with constructs containing 3 to 15 fragments. Constructs with more fragments require more GG sites, making it difficult to distinguish the effects of multiplexing from construct length. To separate these factors, we designed subpools in which we held the total number of GG sites constant (30 per reaction) while varying the fragment number per construct. This meant that reactions assembling shorter constructs contained more total constructs per reaction, whereas reactions assembling longer constructs contained fewer. For example, reactions assembling 3-fragment constructs included 15 independent constructs, while reactions assembling 15-fragment constructs included only two constructs per reaction, with both maintaining a total of 30 GG sites.

We used the OMEGA protocol to assemble designed constructs ranging from 3 to 15 fragments and analyzed assembly fidelity, coverage, and bias using Oxford Nanopore sequencing (**Fig. 2b**). For assemblies with 3 to 12 fragments, over 90% of sequenced constructs were correctly assembled and all target sequences were observed at least once. However, fidelity dropped sharply for constructs with more than 12 fragments. Assembly uniformity remained high up to 8 fragments, with at least 70% of sequences falling within tenfold of the median abundance. Starting at 9 fragments, we observed an increase in over-abundant sequences, though this may be due to the smaller number of total constructs in these reactions, making them more susceptible to random variability. Assessing assembly uniformity becomes challenging beyond 8 fragments due to the low number of constructs in each reaction. As we observed in the multiplexing experiments above, construct abundance does not appear influenced by fragment abundance of the ligation efficiency of the GG sites (**Supplementary Fig. 4 and Supplementary Fig. 5**).

### OMEGA achieves 10x scaling over individual fragment synthesis

OMEGA overcomes the length-versus-scale tradeoff in gene synthesis by combining the low per-base cost of oligonucleotide pools with a simple and efficient assembly process. OMEGA can construct 100s to 1000s of uniquely specified genes depending on the gene length and assembly complexity (**Fig. 3a**). In synthetic biology, metabolic engineering, and protein design, per-gene cost directly impacts the number of constructs that can be built and tested within a fixed budget. OMEGA can substantially reduce the per-gene cost of gene synthesis, expanding the scale of biological engineering.

**Figure 3.**
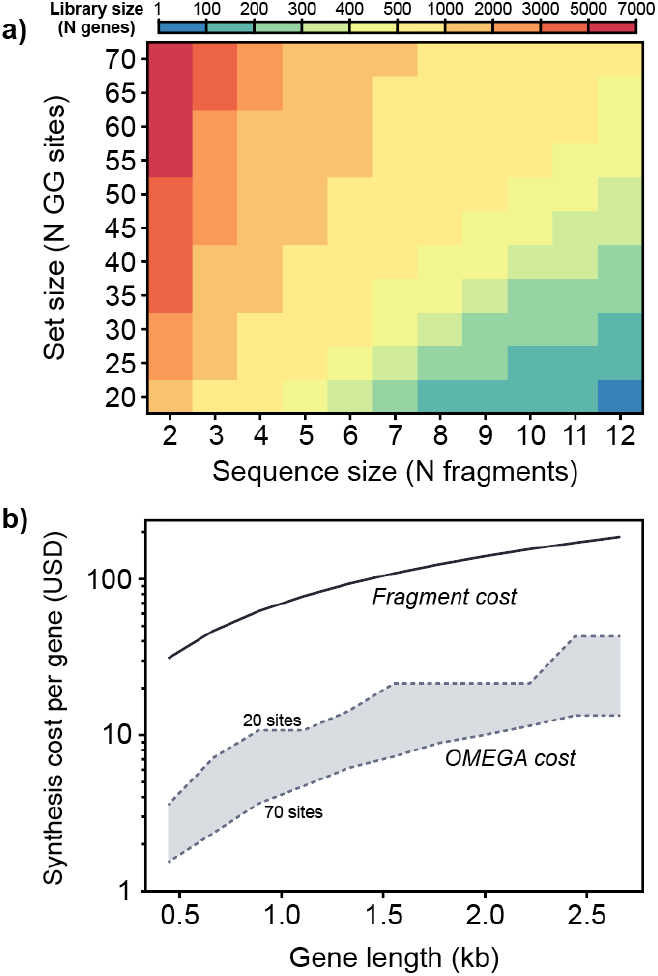
OMEGA assembles hundreds of genes and achieves > 10-fold cost reduction compared to fragment synthesis. **(a)** Library size depends on the number of GG sites, sequence length, and the number of subpools. We use 96 subpools to estimate library size to represent the throughput of a standard 96-well plate. **(b)** Individual costs for sequences (0.3-2.6 kb) synthesized as fragments or assembled from an oligopool using Twist Bioscience pricing. Oligopool gene costs are calculated by dividing the total pool costs by the number of genes in the OMEGA library. We show the range of costs for library sizes using 96 subpools and between 20 and 70 sites per assembly.

To compare the cost efficiency of OMEGA versus individual fragment synthesis, we analyzed Twist Bioscience’s synthesis offerings. Twist provides fragment synthesis up to 5 kb costing 7 to 9 cents per base and fragment synthesis scales nearly linearly with gene length up to the 5 kb limit. OMEGA’s scaling is more complex, as it depends on library size, the fraction of oligos used to encode library sequences (e.g., a 901 bp gene requires four 300 bp oligos, leaving 299 bases unused), and non-linear pricing systems. For example, 2,000 × 300 bp oligos costs 0.6 cents per base while 6,000 × 300 bp oligos cost 0.3 cents per base. To estimate OMEGA’s cost, we modeled assemblies using 20 to 70 GG sites across gene lengths ranging from 0.3 to 2.5 kb. Our analysis showed that OMEGA consistently reduces synthesis costs by at least 10x compared to individual fragment synthesis across this range (**Fig. 3b**).

### Large-scale assembly of natural and synthetic GFP genes

Fluorescent proteins (FPs) have diverse applications as reporters, affinity tags, microscopy, and more.^23^ We have a rich repository of natural FP sequences and the capability to generate synthetic FPs,^24,25^ enabling the discovery of variants with improved spectral properties, brightness, photostability, and maturation times. However, large-scale exploration of FP sequence space remains limited by current gene synthesis capabilities because most sequences are more than 200 amino acids. To evaluate OMEGA’s performance, we tested its ability to synthesize hundreds of full-length FP genes with high accuracy.

We designed a panel of 810 fluorescent protein genes, comprising both natural and synthetic sequences (**Fig. 3a**). We identified 464 Green Fluorescent Protein (GFP) homologs from the UniProt GFP family.^26^ Additionally, we generated a panel of 346 synthetic GFPs (SynGFP) using the generative protein language models (pLMs): CARP-640M, MIF-ST, and ESM-MSA. Sequences were generated through Gibbs sampling and we also included the top double mutants of avGFP predicted by CARP and MIF-ST. Details on the pLMs are provided in the next section. We assessed sequence diversity using a Multiple Sequence Alignment with avGFP and find that sequences vary significantly in both sequence identity and length (**Fig. 3b**). These sequences cannot be generated without custom synthesis and represents a challenging target for OMEGA.

We used the OMEGA software to design an oligopool for assembling our panel of 810 FP genes. Each gene, ranging from 600 to 900 base pairs, was split into three or four fragments for assembly. We distributed the 810 genes across 50 subpools, allowing each subpool to use up to 50 GG sites, two of which were fixed backbone sites. Most subpools contained 16 FP genes per pool. Notable outliers are a UniProt pool that contained 2 genes and a SynGFP pool that contained 4 genes. We followed a high-fidelity assembly protocol developed by Pryor et al.^19^ to construct the FP library in three days with minimal hands-on time. First, we amplified each subpool by performing 50 PCR reactions in microtiter plates, followed by GG assembly within each subpool to construct individual genes. We then combined the subassembly products into separate UniProt-GFP and SynGFP libraries and completed a final column purification. We analyzed the assembled libraries using long-read Pacific Biosciences (PacBio) sequencing at >1,700x coverage to evaluate the assembly coverage, bias, fidelity, and error rates.

We analyzed the sequencing data to determine how many of the designed genes were present after OMEGA assembly (**Fig. 3c**). We only considered perfect sequence matches as correct assemblies. We found 437/464 (94%) of the UniProt-GFP genes and 337/346 (97%) of the SynGPF genes were present at least once. We also found that the assembly protocol resulted in uniform gene abundance, with 88% of the UniProt-GFP genes and 92% of the SynGPF genes falling within tenfold of the median-abundant sequence. Biases in sequence abundance reduce screening efficiency by requiring library oversampling to detect low-abundance sequences. We estimated the screening coverage needed to observe each sequence at least 10 times. We compared our assembled OMEGA libraries to an idealized scenario where all sequences are present in equal abundance (**Fig. 3d**). We found that our libraries generally needed 100X more screening to achieve the same coverage as a uniform library. We also found simulated libraries with 50% incorrect assemblies did not significantly impact screening coverage requirements.

The UniProt-GFP and SynGFP libraries contained overrepresented sequences, prompting us to investigate their cause. While most subpools contained 16 genes, we found that the overrepresented genes originated from remainder subpools with only 2 genes in the UniProt-GFP library and 4 genes in the SynGFP library. This likely occurred because our protocol does not normalize the output from each subpool, and smaller assemblies are more efficient. Biases in sequence abundance were largely attributed to uneven sequence distributions across subpools. To address this, we reassembled the SynGFP library while excluding these remainder subpools, which improved gene uniformity and eliminated overrepresented sequences (**Supplementary Fig. 6 and Supplementary Fig. 7**). Additionally, we found that the number of fragments used per gene (three vs. four) had no impact on gene abundance (**Supplementary Fig. 8**).

The final OMEGA-assembled genes likely contain errors originating from the starting oligopool. Most manufacturers report synthesis error rates between errors per nucleotide, most of which are point mutations.^27^ With a fixed per-base error rate, longer genes are more likely to contain errors and this places an upper limit on the length of error-free genes.^28^ Even when fragments assemble correctly, errors from the original oligos can persist. To estimate the fraction of error-free genes, we mapped OMEGA library reads to the target reference sequences and found over 60% of correctly assembled sequences were error-free (**Supplementary Fig. 9**). This number is consistent with Twist Bioscience’s reported per-base error rate of 1:3000. Additionally, a separate analysis from our GoldenGate multiplexing experiments above showed synthesis errors rarely occurred in Golden Gate or BsaI sites, suggesting that synthesis errors don’t significantly impact assembly fidelity (**Supplementary Fig. 10**).

### OMEGA accelerates generative design and screening of synthetic fluorescent proteins

Generative protein design models can generate vast numbers of sequences with targeted structural and functional properties. Their ability to sample the high-dimensional sequence landscape creates diverse designs with the potential to uncover new and useful proteins.^29,30^ However, bringing these designs to life remains a bottleneck, as synthesizing individual genes is costly, especially for larger proteins. Scalable gene synthesis methods would bridge this gap in the design-synthesis-testing pipeline to accelerate protein discovery.

We tested OMEGA’s ability to assemble genes encoding fluorescent proteins generated by protein language models (PLMs). PLMs learn information-rich protein sequence representations and can be used to direct the generation of new sequences through masked language modeling or next token prediction. We generated GFP variants using three different PLM methods. The first was CARP-640M,^31^ which uses sequence convolutions to learn protein embeddings from aligned related sequences; this embedding is fed into a downstream supervised network that is then trained on GFP sequence-function pairs. The second model was MIF-ST,^32^ a masked inverse folding model that uses a CARP model to learn sequence representations that are fed into a downstream supervised network that is then trained on GFP sequence-function pairs. The final model was ESM-MSA,^33^ a transformer-based protein language model that generates novel protein sequences through masked language modeling from a multiple sequence alignment input.

We generated 100 sequences with CARP-640M, 100 sequences with MIF-ST, and 116 sequences with ESM-MSA. These sequences contained 80-95% mutations from wildtype avGFP. We also used the MIF-ST and CARP-640M models to predict the top 15 avGFP double mutants. As described in the previous section, we applied OMEGA to assemble these 346 SynGFP genes and found 337/346 (97%) were correctly assembled (**Fig. 4c**). This demonstrates OMEGA’s ability to accurately assemble genes with high sequence similarity, highlighting its effectiveness for constructing diverse but closely related variants.

**Figure 4.**
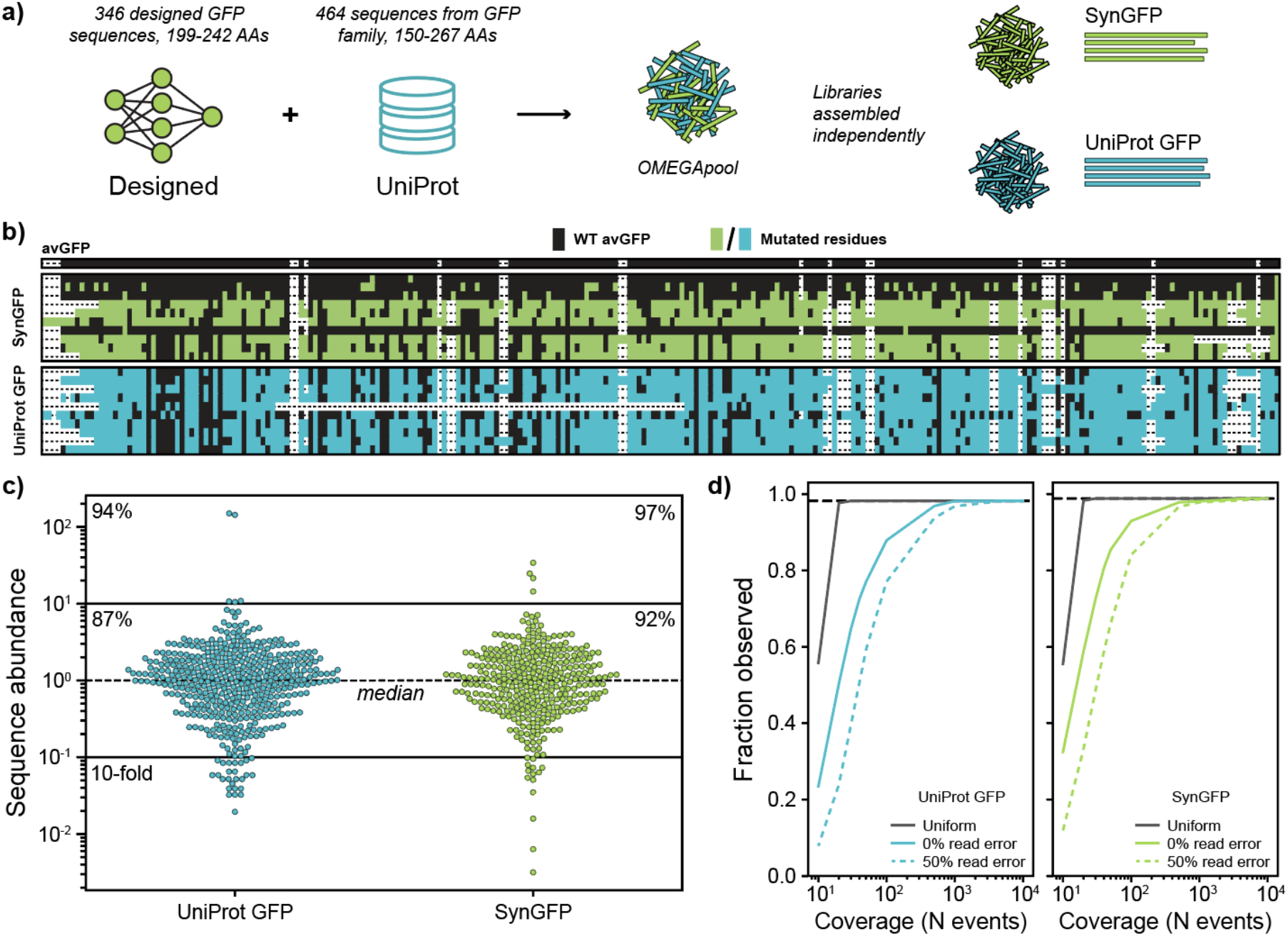
Assembly of 810 fluorescent protein sequences using OMEGA. **(a)** FP sequences were either designed using machine learning (SynGFP) or curated from the UniProt GFP family (UniProt GFP). We used OMEGA to design an oligopool with both SynGFP and UniProt GFP sequences, which we assembled separately using a high-fidelity GG protocol. All subpools used less than 50 GG sites and encoded 2-24 constructs. **(b)** Sequence similarity for a random subset of ten sequences from the SynGFP and UniProt libraries compared to wildtype avGFP. Mutated residues are with respect avGFP. Columns containing >95% gap characters were removed from the alignment. **(c)** Distribution of assembled SynGFP and UniProt GFP sequences. Sequence abundance is represented as x-fold from the median abundant sequence. Percentages within the 10-fold lines indicate fraction of library sequences within 10-fold of the median and percentages in upper corners indicate fraction of sequences observed regardless of abundance. **(d)** Simulated sampling coverage to observe all library sequences. The probability to observe any individual sequence was calculated from the sequence distribution in (c), except for the uniform library, which assumes an equal probability to observe any given sequence. 0% read error indicates that all reads and sequences map to a designed library sequence while 50% read error assumes that 1 in 2 reads will have some error that disqualifies that read. Error does not appear to significantly increase coverage burden.

We evaluated the fluorescence of the assembled SynGFP variants using fluorescence-activated cell sorting (FACS) to sort them into bins based on increasing fluorescence levels **(Fig 5a, Supplementary Table S1)**. To estimate relative brightness, each GFP variant was expressed as a fusion with a constant mKate2 reference protein and a LogF value was calculated based on the variants’ distributions between bins. The screen also included wildtype avGFP as a positive control and incorrectly assembled genes as assumed non-fluorescent negative controls. We fit a Gaussian mixture model to categorize sequences as fluorescent/non-fluorescent (**Supplementary Fig. 11**) and found 26% of all library designs exhibited fluorescence (**Fig. 5b**). We then evaluated each model’s SynGFP design performance (**Fig. 5c**). We found the MIF-ST model successfully designed functional double mutants 80% of the time and was able to identify two variants with a relative brightness higher than wildtype avGFP. The CARP model was less successful with only 35% functional double mutant designs. Using more aggressive MCMC sampling to generate more sequence diversity, both MIF-ST and CARP models fail to design functional fluorescent proteins. We found MCMC sampling of the ESM-MSA model introduced less mutations overall and 61% of generated designs were fluorescent. Two of the sequences designed by ESM-MSA shared 100% sequence identity with known FPs obeYFP and an uncharacterized CFP.

**Figure 5.**
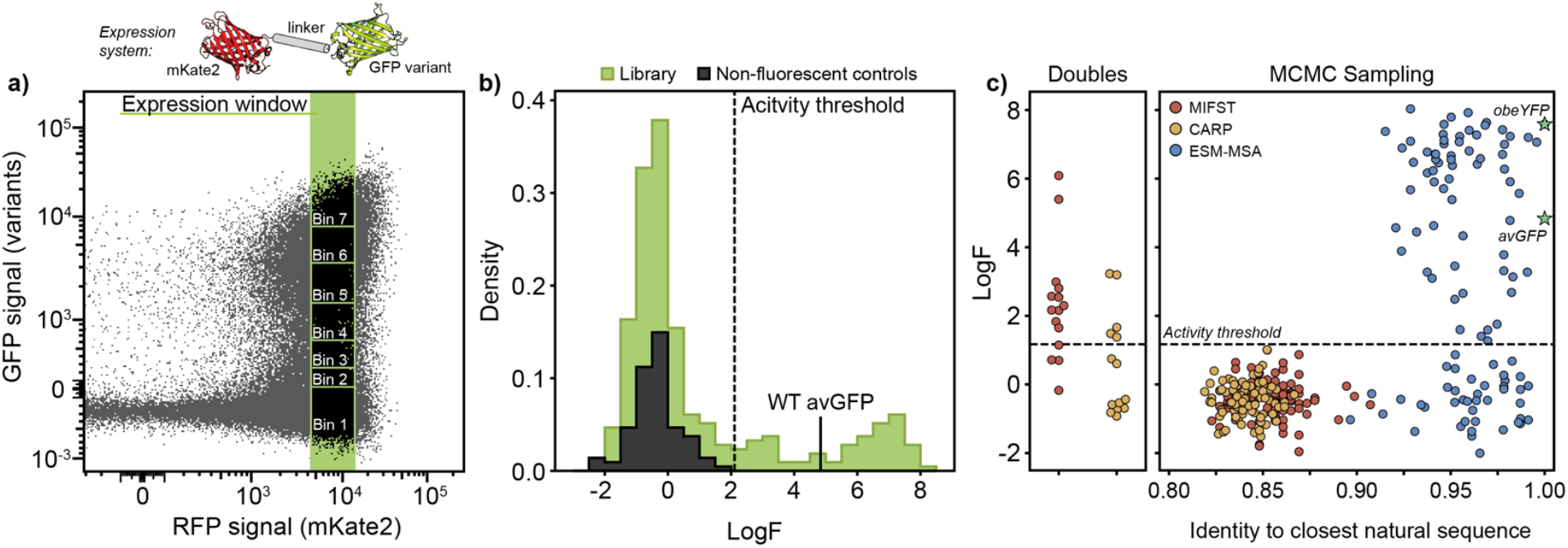
Characterization of SynGFP library using Fluorescence-activated Cell Sorting (FACS). **(a)** Library sequences were expressed as a fusion protein with mKate2, which was kept constant between library variants and indicates level of expression. Cells were sorted into seven bins of GFP fluorescence approximately uniformly distributed across a log-scale within a narrow mKate2 window to control for expression. **(b)** We calculated a log fluorescence (LogF) for each sequence averaged from two replicates. We calculated an activity threshold by fitting observed library sequences with a mixture model to determine the intersection of functional and non-functional populations. We also calculate the LogF for an additional 76 incorrect assemblies that serve as non-fluorescent controls. **(c)** GFP fluorescence plotted by model and sequence identity. Double mutants are 2 mutations from avGFP. Markov chain Monte Carlo (MCMC) sampling sequences are plotted by pairwise-identity to the closest natural sequence in the UniProt GFP family. Two known fluorescent proteins are included as references and indicated by green stars.

## Discussion

Protein engineering relies on large sequence libraries to broadly sample sequence-function space and discover new proteins,^34,35^ but current gene synthesis approaches limit the scale of this exploration. To overcome this, we developed OMEGA, a simple, low-cost, and efficient method for assembling oligopools into large, diverse gene libraries. OMEGA uses a computational pipeline to split genes into short fragments that can be ordered as oligonucleotide pools and assembles them using a hierarchical Golden Gate cloning strategy using Data-optimized Assembly Design (DAD). We demonstrated that OMEGA can reliably assemble constructs of up to 12 fragments and can utilize up to 70 Golden Gate sites while reducing synthesis costs by more than tenfold. To validate the method, we assembled a library of 810 natural and engineered GFP sequences and recovered over 94% of the target sequences. Overall, OMEGA offers a scalable, cost-effective, and robust solution for gene library assembly with broad applications in protein engineering and synthetic biology.

OMEGA leverages standard molecular biology techniques to achieve high-fidelity assembly of large gene libraries through Golden Gate multiplexing. Golden Gate cloning is well-established, with commercially available reagents that enable robust and reliable assembly. Notably, assembling a single OMEGA subpool follows the same straightforward workflow as clonal gene assembly using Golden Gate cloning. DropSynth, another high-throughput gene assembly method, scales Polymerase Cycling Assembly (PCA) by using microemulsions to encapsulate custom-barcoded beads, each carrying the genetic components for individual genes.^16^ This approach requires a complex, multi-step process, including the preparation of custom-barcoded beads and the precise optimization of 20 bp DNA overlaps for each fragment.

OMEGA improves gene recovery compared to existing high-throughput gene assembly methods. DropSynth was used to assemble 346 DHFR genes in duplicate assemblies using different codon sequences. Individual libraries observed 82% and 87% of genes (92% across both assemblies).^36^ In a larger assembly targeting 1536 DHFR constructs, 58% and 68% of genes were observed in two assemblies (∼80% overall). In contrast, OMEGA observes 94-97% of all target sequences in a single assembly with libraries of similar or larger size. The SynGFP and UniProt GFP can be considered as a single large library since all sequences were ordered in a single oligopool and assembled in the same way. In total, we observe 774/810 genes (95%) of all GFP constructs, suggesting that library coverage does not significantly decrease with library size. Furthermore, OMEGA maintains high fidelity for constructs with up to 12 fragments while DropSynth assembly fidelity significantly drops around ∼5 fragments. We expect that the easy integration of OMEGA assemblies into standard Golden Gate cloning workflows, combined with our high fraction of observed target sequences, will enable scientists to construct gene libraries with ease.

Innovation in synthetic biology is driven by our capacity to design and synthesize custom DNA. Golden Gate cloning is the foundation of many cloning workflows to efficiently design DNA sequences using multifragment assemblies. Data-optimized Assembly Design (DAD) uses experimentally derived overhang annealing and ligation preferences to identify high-fidelity Golden Gate cloning junctions, significantly expanding the scope of GG cloning.^19^ Here, we extend the complexity of DAD assemblies from 52 GG sites to 70 in a single reaction, utilizing ∼30% of all possible 4-bp overhangs and achieving ∼95% assembly fidelity. We also demonstrate that assemblies with up to 12 fragments maintain high fidelity, underscoring DAD’s potential for constructing even larger, high-complexity assemblies beyond protein-coding genes. The success of our assemblies, which each use unique combinations of GG sites, emphasizes that DAD is a robust method to design novel sets of GG sites that yield high-fidelity assemblies, reducing constraints based on sequence and assembly size.

We expect OMEGA’s capabilities will improve alongside advances in DNA synthesis technology. Its primary limitations stem from synthesis constraints and DNA binding energetics. Oligo synthesis errors impact library quality by reducing the fraction of error-free sequences setting an upper limit on gene length.^28^ While future improvements in synthesis accuracy will help overcome these challenges, existing methods such as MOPPS or enzymatic error correction can already mitigate errors.^27,37^ The number of GG sites per reaction is constrained by the number of orthogonal fixed-length overhang interactions and limited by the capabilities of Type IIS restriction enzymes. Improvements to DNA synthesis technology will naturally extend OMEGA’s assembly length and fidelity by providing longer fragments with fewer errors. For example, Twist Bioscience recently introduced multiplexed gene fragments up to 500 bp, which would enable OMEGA to assemble constructs up to 5 kb. With continued improvements in synthesis capabilities, OMEGA could support multikilobase assemblies, expanding its applications to metabolic engineering, synthetic plasmid construction, and large-scale protein design.

We are in an information age of biological data where it is increasingly important to target sequence space with precision-designed libraries to develop and refine ML engineering strategies. OMEGA is a simple gene assembly method that excels in assembling libraries of sequence-diverse constructs with high coverage from oligopools. With Golden Gate cloning, OMEGA libraries integrate seamlessly into established cloning workflows and mitigates financial barriers that limit the scope of custom libraries. We envision that OMEGA will empower data-driven engineering with its simple and affordable library workflow.

## Methods

### Implementation of OMEGA software

The OMEGA software automates library design to assemble diverse genetic constructs from oligopools. Users provide a list of codon-optimized sequences and OMEGA returns two files detailing the precise oligo sequences to order and a list detailing the contents of each subpool. The main function of the OMEGA software is to use DAD to find fragmentation patterns with optimal GG sites to enable high-fidelity assembly of individual genes. We calculate the fidelity as described in Pryor et al.^19^ that assumes each GG site will be used in a single sequential assembly. The overall fidelity is thus the product of the individual site’s predicted fidelity, which estimates the likelihood of a correct ligation in the presence of other GG sites in the set.

To start, the user provides a list of codon-optimized sequences and specifies the Type IIS restriction enzyme and the maximum number of GG sites that should be used for library assembly (GG site budget). If needed, the user may also provide two backbone ligation sites that are included in the fidelity prediction and set size calculation. OMEGA begins by estimating the number of fragments needed to encode the longest gene in the library and uses this value as the target number of fragments to break all genes into. Breaking genes into the same number of fragments reduces potential library bias from shorter genes that are more efficiently assembled than longer genes. OMEGA calculates how many pools are required to encode all genes based on gene length and GG site budget and uniformly distributes genes amongst the pools such that the pool with the most genes has, at maximum, one more gene than the pool with the fewest genes.

Next, OMEGA computationally designs fragments for each subpool by optimizing the fidelity of the GG sites created by different fragmentation patterns. OMEGA initializes pools by breaking genes into fragments of equal length. Any sites that are duplicated are shifted left or right by 1 bp until all starting sites are unique within a subpool. The challenge of fragment design can be posed as a combinatorial optimization problem guided by the predicted fidelity of the current set of GG sites. Each gene can be broken into one of several legal fragmentation patterns that yield different sets of GG sites. This problem space is too large to solve absolutely – for example, a pool with 10 genes that each have 3 break points and each break point has 25 GG site options has ∼10^41^ site combinations (including combinations with repeated GG sites).

We apply simulated annealing to approximate an optimal solution from these potential options. For every optimization step, we shuffle a random GG site to propose a new set of GG sites that differ by one position. If the predicted fidelity is better than the previous solution, the new site is accepted. If the solution is worse, then it is accepted with probability *e*^*ΔFidelity/T*^ where *T* is decreased along a logarithmic temperature gradient ranging from 5 × 10 ^−3,^ to 10 ^−5^ over 2,000 to 10,000 optimization steps. This optimization process is completed *N* number of times using different random seeds to find different solutions. The best of N optimization runs is selected for subpool design. More optimization steps and optimization runs are recommended for assemblies using more GG sites. OMEGA automatically packages genes across all subpools into a list of fragmented sequences ready to be ordered with all required restriction enzyme sites and amplification primers added in. All oligo fragments are padded with random DNA sequences between the BsaI binding site and primer binding site to make oligos a uniform length. An additional list is provided that provides details on each pool, including library genes, predicted fidelity, GG sites, primers for pool amplification, and the random seed used to design the gene fragments.

We provide OMEGA as the ggopt python package and provide a conda environment with all necessary packages to run it. We use jsonargparse to make customizing OMEGA’s runtime parameters to suit specific use cases. For example, users may change the restriction enzyme, number of GG sites used in each subpool, backbone sites, and more. We provide an example configuration file with all parameters users can change in the SI and provide examples of library optimizations in the code published with this paper.

### Design of diverse Golden Gate sets

We used OMEGA to design high-fidelity, diverse GG sets containing between 20 and 70 sites in increments of five. We required that all sets contain the vector ligation sites AATG and TTAG, but otherwise allowed the use of any GG sites. We designed 50 sets for each size and identified the two most diverse sets using pairwise Hamming Distance as a diversity metric for a total of 22 unique Golden Gate sets (11 sizes, 2 replicates). Sets are referred to by their size and replicate (ex. sets 30a and 30b refer to our two diverse sets containing 30 sites). All 22 sets are used in the 2-fragment assemblies that vary GG site number. The assemblies testing the effect of fragment number use sets 30a and 30b for all fragment numbers.

### Design of OMEGA parameterization assembly sequences

We designed synthetic sequences for assemblies varying GG site number and assemblies varying fragment number used to parameterize OMEGA. The synthetic sequences consist of a series of unique 25mer barcodes joined by a variable GG site that are not part of the barcode sequence. Barcode sequences were chosen from a set of super orthogonal barcode sequences designed by Xu et al.^38^ All barcodes vary by at least 15 bp to allow for good discrimination of sequences using both Illumina and Oxford Nanopore sequencing platforms and do not contain any BsaI binding sites. The first and last fragments are flanked by the upstream and downstream vector ligation sites AATG and TTAG.

For each of the 22 unique sets of GG sites, we designed 2-fragment assemblies using the above synthetic sequence design scheme. For the assemblies where we varied the number of fragments, we use sets 30a and 30b to randomly assign the GG sites connecting between 3 and 15 fragment assemblies. For each fragment number, we designed as many sequences as possible given a fixed GG site budget of 30.

To design the genes as oligos, we fragmented sequences by barcode and added corresponding upstream and downstream GG sites, BsaI restriction sites, and primer binding sites. We added equal padding to the left and right of the barcode between the BsaI restriction site and primer binding site to ensure that all sequences were 120 nt. We provide a full list of genes, sequences, GG sites, and primers in the supplementary information.

### GFP sequence design

For the SynGFP library, we designed 346 GFP sequences using different ML models. First, we used MIF-ST and CARP-640M to score every possible double mutant to wild-type GFP and selected the top 10 from each model. Next, we performed Gibbs sampling using MIF-ST, CARP-640M, and ESM-MSA, as described in [39]^39^, [40]^40^ to diversify natural GFP sequences. Beginning with a natural sequence, at each iteration, 10% of positions were masked, and the model was used to predict the probability of each amino acid at each masked position. For each of 10 burn-in steps, the masked positions were sampled from the predicted distributions. Then, for each of 10 sampling steps, each masked position was set to its most probable amino acid, resulting in 10 generated sequences per starting natural sequence. For CARP-640M, 10 starting sequences were randomly chosen from a set of putative GFP homologs. For MIF-ST, samples used the PDB 2WUR structure and corresponding sequence. For ESM-MSA, 10 subsets of 63 aligned homologs and one starting sequence were randomly chosen as starting points. This results in 100 generations for each model.

For the UniProt library, we randomly selected 464 sequences ranging between 200 and 250 amino acids from the UniProt GFP family (accessed: 07-19-2022).

All sequences were codon optimized for yeast using the GenSmart codon optimization tool from Genscript and specified that optimized sequences excluded BsaI sites. We used OMEGA to design the SynGFP and UniProt GFP libraries separately, though the same design parameters were used. In both cases, we specified that no more than 50 GG sites could be used in each subpool and that each pool must include AATG and TTAG as vector ligation sites. Subassembly pool sizes ranged from 20 to 4 genes for the SynGFP library and 24 to 2 genes for the UniProt library. All genes were broken into either 3 or 4 fragments. We ordered the OMEGA library as 300-bp oligos and added random DNA padding between the primer amplification site and BsaI restriction site to ensure all oligos were 300 bp. Subassembly pools are indexed by a unique combination of orthogonal primers for selective amplification.^22^ For a full list of primers and sequences, please refer to the supplementary information.

### High-fidelity Golden Gate assemblies

All OMEGA assemblies in this work follow the same PCR amplification and assembly protocol. Oligopools were resuspended to a concentration of 1 ng/uL. Subassembly pools were amplified directly from the stock oligopool using Kapa HiFi HotStart ReadyMix using 1 ng of template DNA (95 ºC 3 mins → (98 ºC 20 secs → 61 ºC 15 secs → 72 ºC 15 secs) × 35 cycles → 72 ºC 1 min) and cleaned up using Zymogen DNA Clean and Concentrate (Zymo Research, #D4006). Pool-specific primers are provided in the supplementary data.

Golden Gate assemblies were performed for each subpool in parallel on a 96-well plate following highfidelity assembly protocol described in Pryor et al.^19^ First, insert and destination cassette are digested with BsaI- HFv2 with an 18:1 molar ratio for two hours at 37 ºC (15u BsaI, 1X T4 Ligase Buffer, 20 uL reaction volume). Following incubation, 1,000 U of T4 Ligase was added directly to the digest and incubated for at least 18 hours (37 Cº) followed by a final incubation to neutralize BsaI (65 ºC, 15 mins). All assemblies were cleaned up using Zymogen DNA Clean and Concentrate. Subassemblies from the same library were pooled before cleanup.

Sequences from the OMEGA parameterization assemblies varying GG set size and fragment number were assembled into a linear destination cassette that we designed and ordered as a gene fragment from Twist Bioscience. We designed reverse-oriented BsaI binding sites separated by five using CTAA for the upstream site and CATT for the downstream site. There is a randomized 10-bp upstream of the first BsaI site. Primers flank both GG sites for subsequent PCR after assembly. The GG product from each of these assemblies was amplified using PCR (95 ºC 3 mins → (98 ºC 20 secs → 61 ºC 15 secs → 72 ºC 15 secs) × 16 cycles → 72 ºC 1 min; primers: pr.subra_84, pr.subra_88), digested with BsaI-HV2 for 30 mins. at 37 ºC (15u, 1X CutSmart buffer), and cleaned up using Zymogen Clean and Concentrate. Product sizes were analyzed on Tapestation.

The UniProt and SynGFP libraries were assembled into a modified pqe30 expression vector previously described.^5^ We started from a related backbone (Addgene, #74748),^41^ which expresses a KillerOrange-GFP fusion protein. We replaced KillerOrange with an mKate2 construct ordered from Twist Bioscience using the BamHI and NotI restriction sites. We next replaced GFP with the BsaI sites AATG and TTAG using PCR and blunt-end ligation (pr.001 and pr.002). We also created a version without mKate2 using PCR to remove the mKate2 sequence followed by blunt end ligation (primers: pr.003, pr.004). We assembled the SynGFP library into the vector with mKate2 and UniProt GFP library into the vector without mKate2. We also performed a second assembly of the SynGFP sequences called SynGFP B, which excludes two pools from the SynGFP assembly that included 3-fragment genes or had fewer genes than other subpools. A list of all subpools and sequences used in each assembly is provided in the supplementary materials.

### NGS sequence processing for parameterization assemblies using variable GG sites

We applied 2×150 Illumina sequencing to all GG assemblies. 2×150 sequencing can read sequences up to ∼300 bp. Correct 2-fragment assemblies are 62 bp, but assemblies with up 8 fragments are short enough for 2×150 reads. We expect that 2×150 sequencing can adequately detect assembly results. All subpools were sequenced with a minimum of 26,000X coverage. The full data processing pipeline is included in the supplementary materials. Briefly, we merged forward and reverse reads, trimmed extraneous DNA outside of the insert region (30 bp from start, 20 bp from end), and discarded reads with a predicted error rate > 1 bp. We then mapped individual barcode sequences against reads to define an assembly code, which is determined by the order in which the barcodes were assembled and identifies assemblies independent of sequence. Since oligopools contain synthesis errors and our objective is to assess the effect of GG set size on assembly fidelity, we allow up to 3 mismatches in the barcode sequence. We discarded any partially matched reads, any assembly that contained barcodes from more than one subpool, and if the trimmed read did not start and end with the expected flanking GG sites.

We use the assembly code to differentiate correct and incorrect assemblies. All counts were aggregated by assembly code to assess GG site fidelity independent of synthesis errors in the barcode sequence. Fully mapped reads define the full read population and correct assemblies are those with predicted assembly codes. Target sequences were considered assembled if the corresponding assembly code was observed at least once.

### NGS sequence processing for parameterization assemblies using variable fragment numbers

We applied Oxford Nanopore Sequencing to all GG assemblies. The expected size of assemblies is between 141 bp and 489 bp and some incorrect assemblies may be longer. Oxford Nanopore sequencing is less biased by variable length sequencing input and can sequence longer constructs than Illumina. All subpools were sequenced with a minimum of 35,000X coverage. The full data processing pipeline is included in the supplementary materials. Briefly, we used usearch to map individual barcode sequences and flanking primer sequences to ONT reads, allowing for up to 3 mismatches. We kept reads that satisfied the following conditions: 1) both upstream and downstream primers were present once and in the same direction, 2) aligned barcodes covered the whole insert region, and 3) all barcodes were from a single assembly pool. For all analyses, we considered the total number of reads as those with full-coverage alignments and barcodes from the same assembly pool. We used the order of barcode assembly to assign each read an assembly code and consolidated counts by assembly code. Correct reads are those with predicted barcodes. Target sequences were considered assembled if the corresponding assembly code was observed at least once.

### Sequencing for oligopool

To assess the quality of amplified oligopools, we applied 2×150 Illumina sequencing to the amplified oligos for each subpool used in the OMEGA parameterization assemblies. To process the data, we used the 32-bit version of usearch.^42^ We include the full data processing pipeline in the supplementary materials. Briefly, we merged forward and reverse reads and filtered out any reads with a predicted error rate greater >1 base. We counted the frequency of each unique sequence and discarded any sequence with < 10 counts.

We generated two alignments from the oligopool data. To assess synthesis errors across the full oligo sequence, we used the ‘-search_oligodb’ command to map the designed oligo sequence to Illumina reads and allowed up to 3 mismatches. Only reads that were 120 bp were kept. Correct sequences were those with perfect matches to designed oligos and incorrect sequences contained at least one error. To assess synthesis errors in the BsaI binding site and GG site, we mapped insert sequences beginning and ending with the first and last bp of the BsaI binding sites and allowed up to 3 mismatches and only kept reads that were the expected 47 bp. We counted the frequency of errors in the BsaI and GG site.

### PacBio sequencing for UniProt GFP and SynGFP Golden Gate assemblies

We applied long read PacBio sequencing to the SynGFP, SynGFP B and UniProt GFP assemblies to assess assembly fidelity. For each library, the assembly product was transformed into NEBExpress I^q^ competent cells (New England Biolabs, #3037I) and the full transformation, minus a small volume to measure transformation efficiency, was plated onto LB+carb plates. Plates were incubated overnight at 37 ºC and then moved to 4 ºC for another 24 hours. Aft this, an equal fraction of cells from each plate was collected and added to 50 mL growth media (25 g/L Luria Broth, 10 g/L molecular biology grade agar, 1% glucose) and grown until ∼1OD. We harvested the DNA using a plasmid prep kit (QIAGEN, #12943), linearized the DNA using NotI-HF (New England Biolabs, #R3189) following manufacturer protocol, and submitted samples to the University of Wisconsin – Madison Biotechnology Center. Sequencing achieved at least 16X coverage based on the number of unique transformants.

We used ‘pbmm2’ to map library sequences to full-length PacBio reads. Library sequences included ∼25 bp upstream and downstream flanks of the destination vector sequence to ensure assembled sequences and plasmid were contiguous. We kept reads that satisfied the following criteria: 1) library sequence was contiguous with the destination vector, 2) the full read length was within 200 bp of the predicted read length. We categorized mapped reads as correct if the sequence perfectly matched the designed sequence and considered sequences assembled if a perfect sequence match was present at least once. Sequences with up to 5 mismatches and 0 insertions and deletions were used for additional synthesis error analysis.

We further identified a subset of incorrect assemblies in SynGFP that we use as non-fluorescent controls in our characterization of the SynGFP library. We looked for sequences between the vector ligation sites that occurred more than 100 times that did not match any known library sequences. We selected a subset of 116 of these to create a set of negative sequences as non-fluorescent controls.

### Experimental evaluation of SynGFP sequences

We measured GFP fluorescence for the SynGFP library using fluorescence-activated cell sorting (FACS) as previously described by Sarkysian et al.^5^ Briefly, the SynGFP library was transformed into NEBExpress I^q^ competent cells (New England Biolabs, #3037I) following manufacturer’s protocol and plated on Luria Broth-carbenicillin (LB-carb) agar plates (25 g/L Luria Broth, 10 g/L molecular grade agar). Plates were incubated overnight at 37 Cº and then incubated for 24 hrs at 4 Cº to allow for expression and fluorescence maturation. Plates were washed with 4 mL LB+carb (25 g/L Luria Broth, 100 uM carbenicillin) and used to create 1 OD expression cultures. Duplicate cultures were induced with 1 mM isopropyl *β*-D-1-thiogalactopyranoside (IPTG) and incubated for 2 hrs (23 Cº, 230 RPM). We spiked in wildtype avGFP such that it was 1% of the total library. After incubation, 1 OD of cells were spun down and resuspended in ice cold phosphate buffer (1X PBS).

We sorted GFP variants based on fluorescence intensity. We used ex561/em670 and ex488/em530 wavelengths to detect mKate2 and GFP fluorescence, respectively. Compensation was applied to remove overlapping signal. To account for variations in expression, we first defined a narrow gate for mKate2 expression and then defined a series of 7 sort gates for GFP expression within this population. The GFP gates followed an approximately uniform distribution across the logarithmic scale with slightly larger gates for the least and most fluorescent populations. Sorted cell populations were grown to 0.6-1.0 OD in LB-Carb media with 1% glucose to suppress protein expression and reduce growth bias due to protein expression. The DNA was harvested from 50 mL cultures using a QIAGEN Plasmid *Plus* Midi kit (Qiagen, #12943).

Sorted FACS populations were sequenced using Oxford Nanopore and used to calculate a LogF value for all SynGFP sequences and the incorrect assemblies. We used ‘pbmm2’ to map reads to library sequences. The DNA was linearized using tagmentation, which may cleave a single plasmid at multiple points. We could not use read length to filter reads as with our PacBio samples and instead only required that mapped library sequences were contiguous with the destination vector and overlapped by at least 25 bp. Library sequences were very diverse with the closest related sequences sharing ∼90% sequence identity. Assembly fragments also varied in size, which would create significant size disparity in the event of incorrect assemblies. We reasoned that incorrect sequences would vary significantly from designed sequences and allowed mapped reads to have up to 5 mismatches and 6 insertions and deletions.

Finally, we filtered sequences for synthesis errors by removing ONT sequences with known or novel point mutations at positions where we observed synthesis errors in the PacBio data. For each sequence, we aligned mapped ONT reads with the mapped PacBio reads described previously. Synthesis errors were identified as positions where at least one read reported a point mutation with a qscore >= 80. We removed any reads with incorrect bases at these positions, but otherwise permitted some errors in other portions of the sequence. The PacBio sequencing was greater library coverage than the Oxford Nanopore sequencing, and we reasoned that Oxford Nanopore sequencing would not observe any novel point mutations.

We calculated a log fluorescence value for each sequence based on its probability distribution across GFP bins following previously described methods.^43^ We excluded sequences from analyses if they had fewer than 55 combined counts across bins in either replicate. We report LogF values as an average of replicate LogF scores since there was a strong correlation between replicates (Pearson’s *ρ*: 0.94).

## Notes

### Competing Interest Statement

The authors have declared no competing interest.

### Summary of Updates

We have included links to the software repository.

### Funding

- National Science Foundation: 2226383

