## Supplementary Information for "Scalable and cost-efficient custom gene library assembly from oligopools"

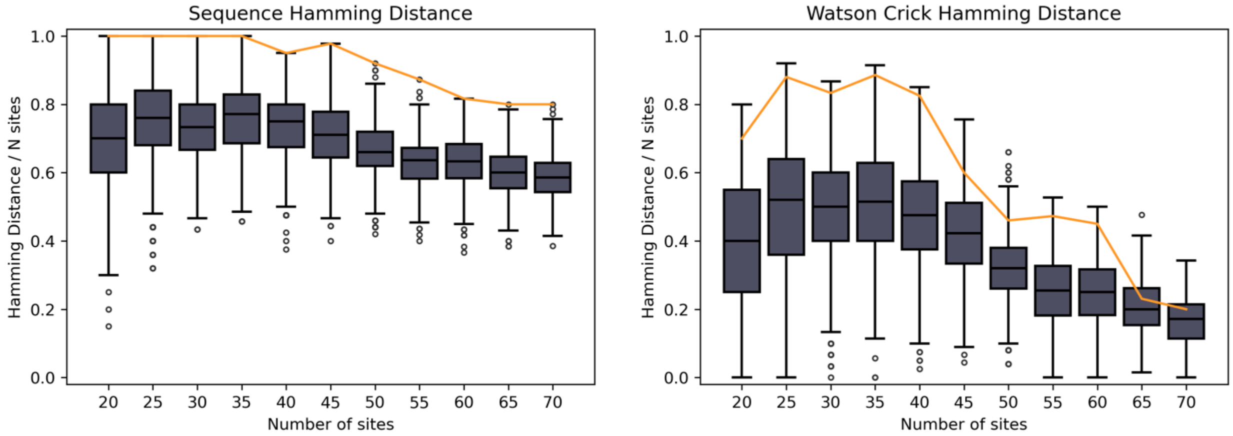


**Supplementary Figure 1. Pairwise hamming distance of optimized Golden Gate sets using 20 to 70 sites.** Boxplot shows pairwise HDs for all designed sets, orange line shows the pairwise HD for the two experimentally validated sets. **(a)** shows diversity in terms of sequence identity and **(b)** shows diversity in terms of Watson Crick pair.

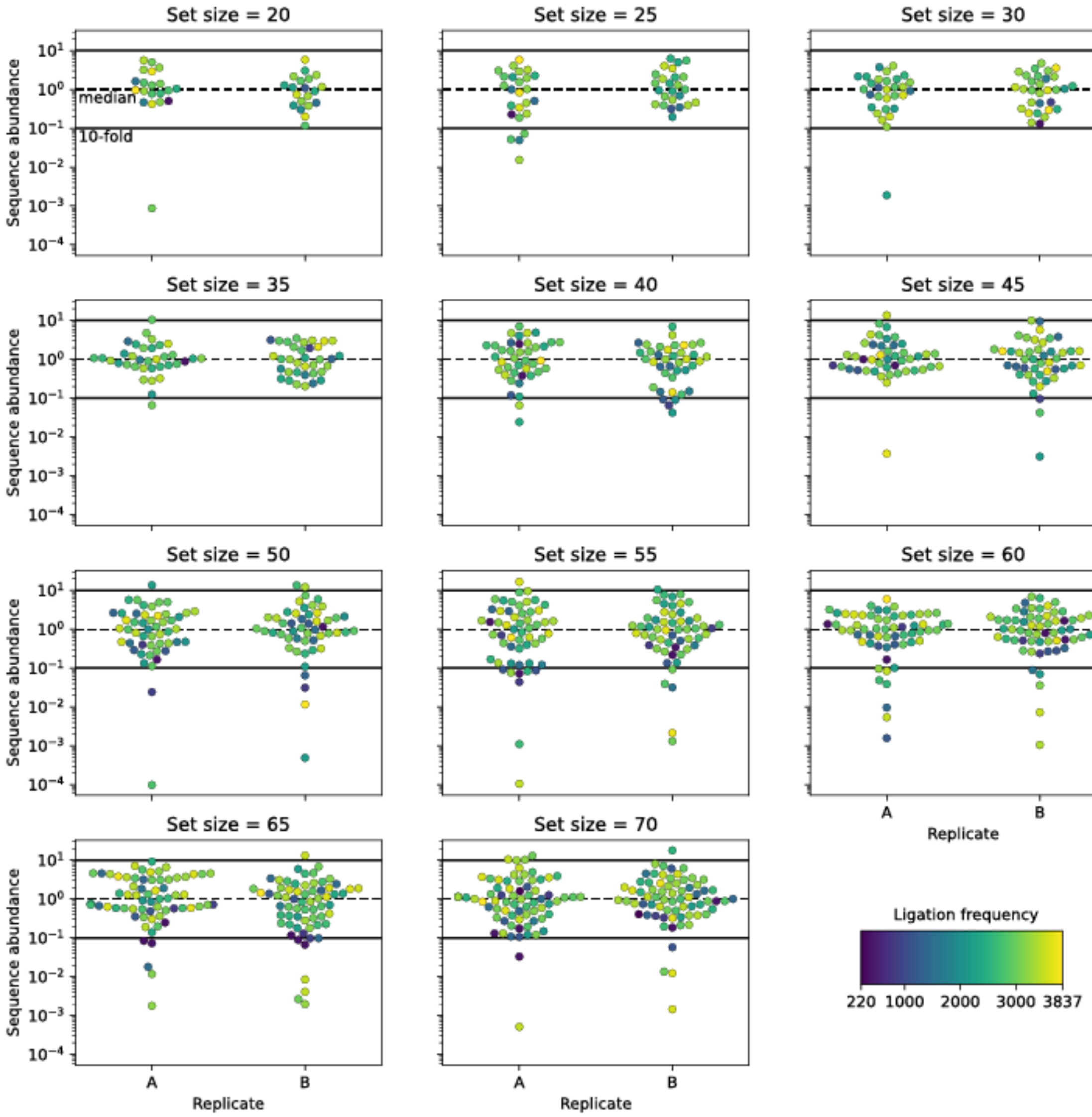


**Supplementary Figure 2. Correlation between sequence abundance and ligation frequency for assemblies using 20 to 70 Golden gate sites.** Potapov et al.[1] estimate ligation efficiencies of individual Golden Gate sites as the number of correct ligation events in their ligation frequency experiments. High and low ligation frequencies correspond to high and low ligation efficiencies. Here, assembled sequences are plotted for all replicates for each set size according to the x-fold sequence abundance. The hue corresponds to ligation frequency for of the designed GG site connecting the two fragments. Low-efficiency sites are used in sequences with both low and high-abundance with no noticeable trend. In large assemblies, very low ligation efficiency may minorly affect sequence abundance.

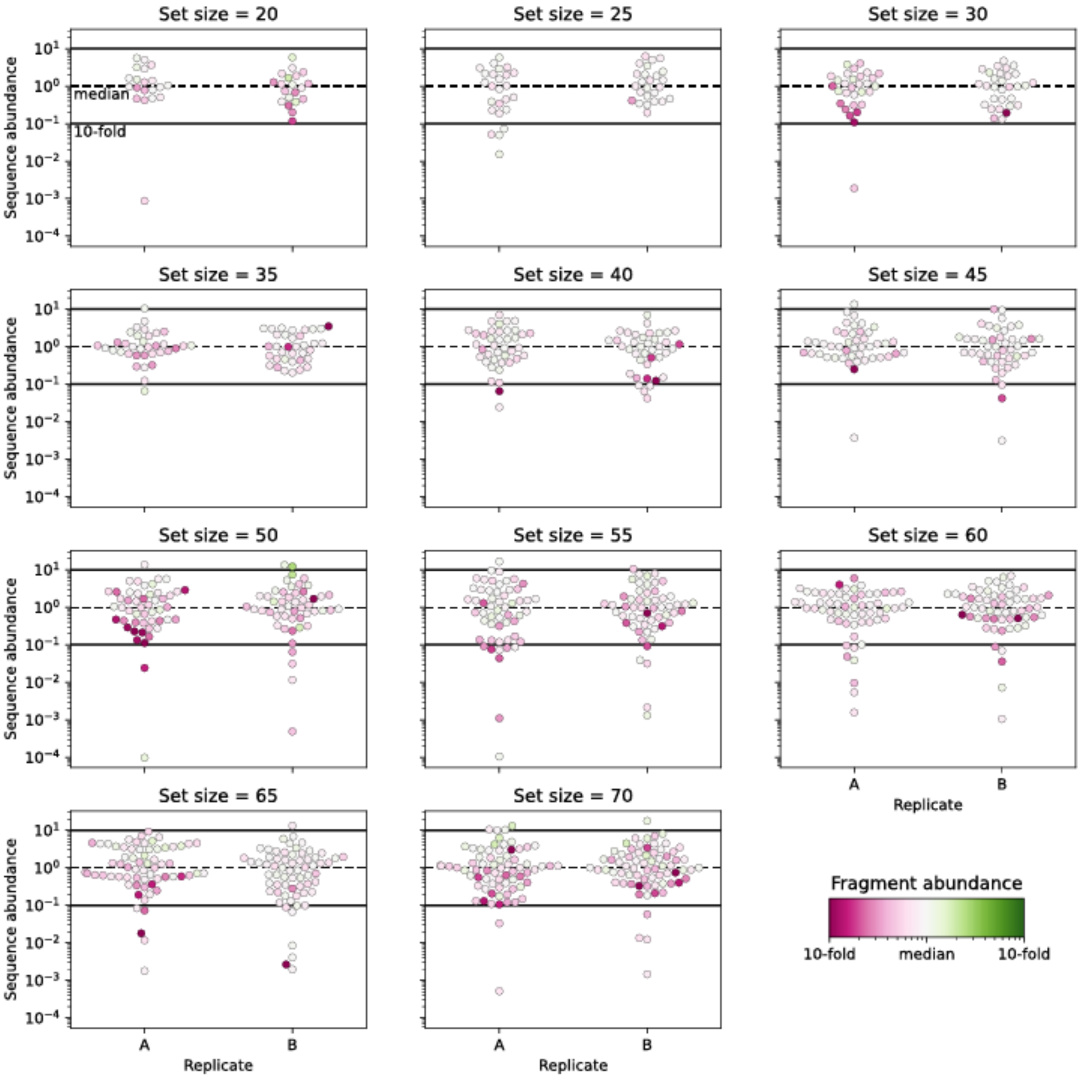


**Supplementary Figure 3. Effect of fragment abundance on sequence abundance for assemblies using 20 to 70 Golden gate sites.** We sequenced the amplified oligos used as input to the GG assembly and calculated the relative abundance of each fragment with respect to the median abundant fragment for each assembly pool. Here, assembled sequences are plotted for all replicates for each set size according to the x-fold sequence abundance. The hue corresponds to the least abundant fragment used in the assembly. In larger assemblies using more than 45 sites, assemblies with low-abundant fragments tend to result in low-abundant assemblies.

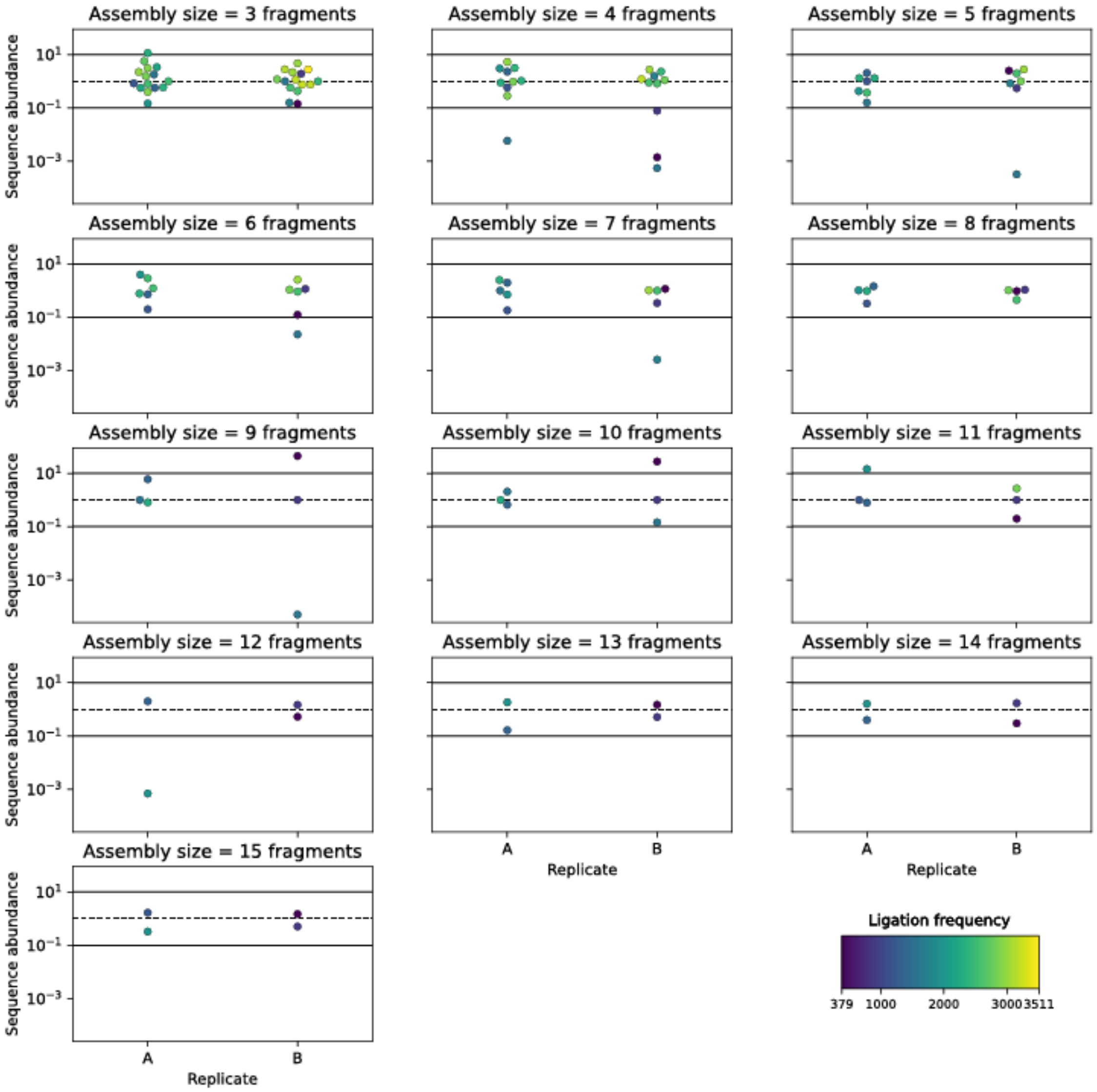


**Supplementary Figure 4. Correlation between sequence abundance and minimum ligation frequency for assemblies using 3 to 15 fragments.** Potapov et al.[1] estimate ligation efficiencies of individual Golden Gate sites as the number of correct ligation events in their ligation frequency experiments. High and low ligation frequencies correspond to high and low ligation efficiencies. Here, assembled sequences are plotted for all replicates for fragment number according to the x-fold sequence abundance. The hue corresponds to ligation frequency for of the lowest efficiency GG site. Low-efficiency sites may result in low-abundance assemblies.

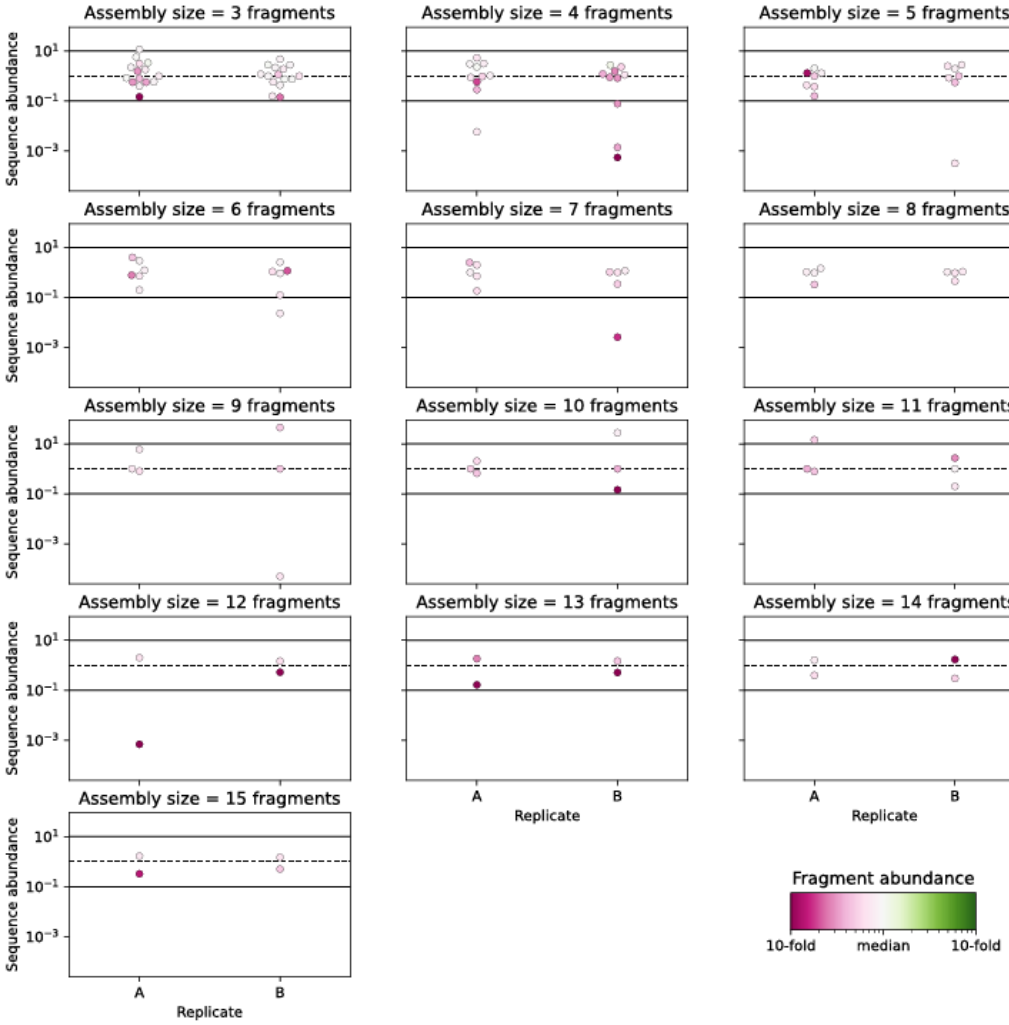


**Supplementary Figure 5. Effect of fragment abundance on sequence abundance for assemblies using 3 to 15 fragments.** We sequenced the amplified oligos used as input to the GG assembly and calculated the relative abundance of each fragment with respect to the median abundant fragment for each assembly pool. Here, assembled sequences are plotted for all replicates for each fragment number according to the x-fold sequence abundance. The hue corresponds to the least abundant fragment used in the assembly. Low-abundant fragments appear to be slightly more common in low-abundant sequences.

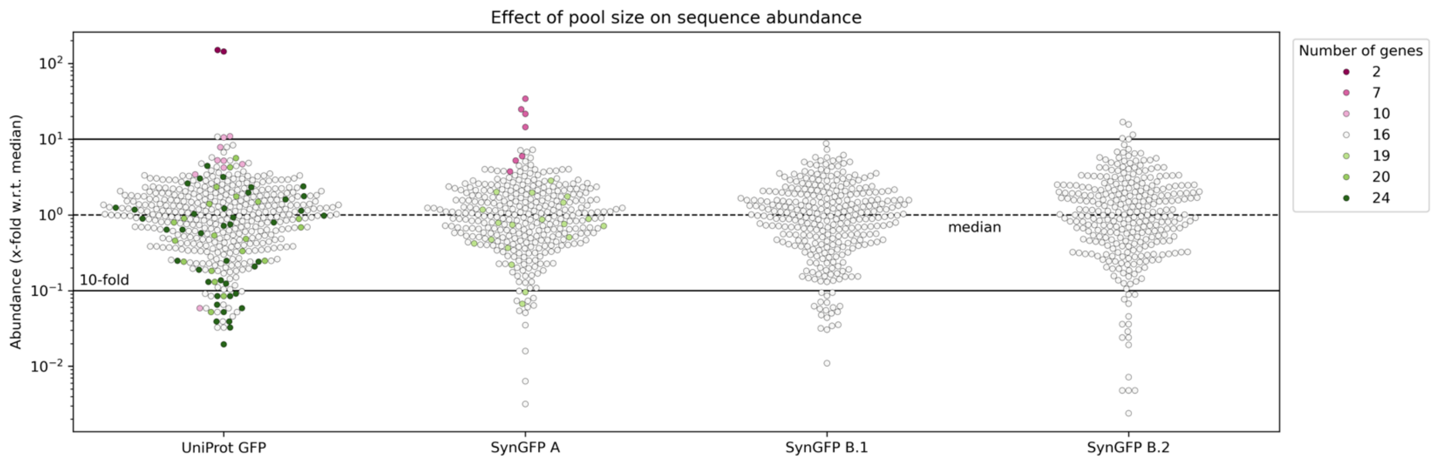


**Supplementary Figure 6. Varying the number of sequences in sub-assembly pools biases library sequence distribution.** Library assemblies were sequenced with long read PacBio sequencing and plotted as the x-fold with respect to the median-abundant sequence for each library. Color indicates the size of subassembly pools (ex. white indicates that sequence was assembled with 15 other sequences for a total pool size of 16). Sequences from small assembly pools (<16) are more abundant than larger assemblies. Large assemblies (>=20) may result in under-represented sequences.

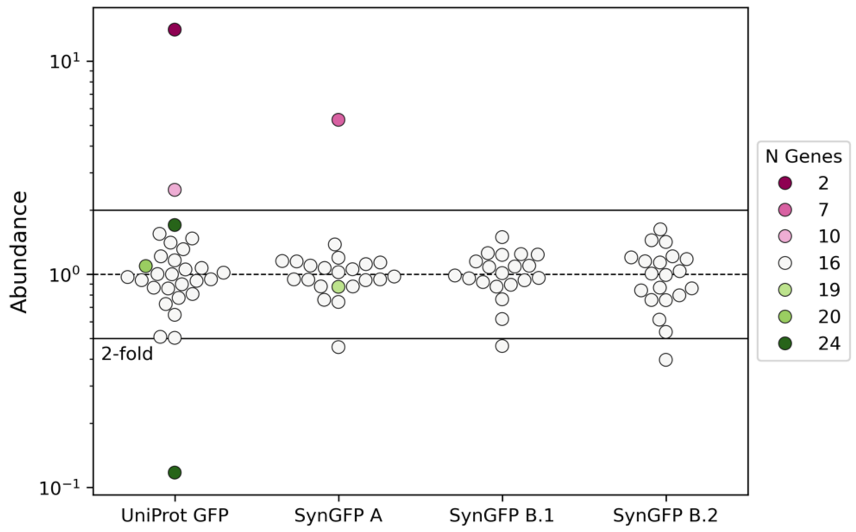


**Supplementary Figure 7. Assembly uniformity across subassembly pools.** We assessed if all sub-assembly pools were equally represented in the combined library. We combined assembly products from all pools for each library without normalization and applied long read PacBio sequencing. The total number of reads were counted for each pool and divided by the median-abundant sub-assembly pool to represent pool abundance as x-fold from the median. Sub-assembly pools are colored by the number of genes assembled in the pool. Most pools assemble within 2-fold of the median. Pools that assemble fewer genes are typically over-abundant in the library (UniProt GFP and SynGFP A). Removing pools with fewer genes mitigates assembly bias (SynGFP A vs. SynGFP B).

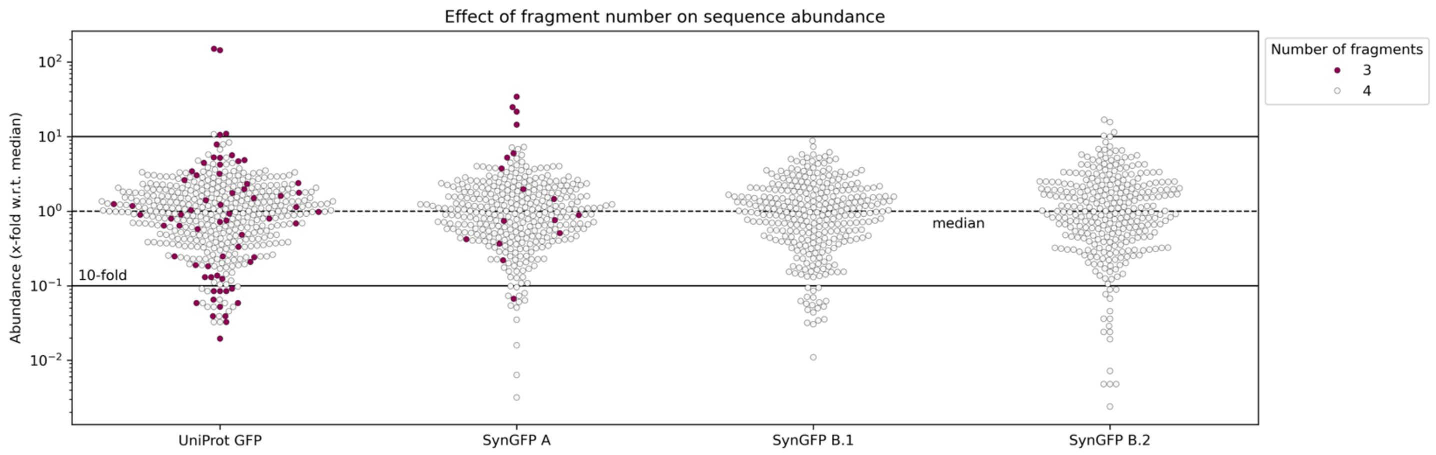


**Supplementary Figure 8. Small variation in sequence length does not bias library assembly.** Library assemblies were sequenced with long read PacBio sequencing and plotted as the x-fold with respect to the median-abundant sequence for each library. Color indicates the number of fragments used to assemble sequences. Three-fragment assemblies are present across all sequence abundances.

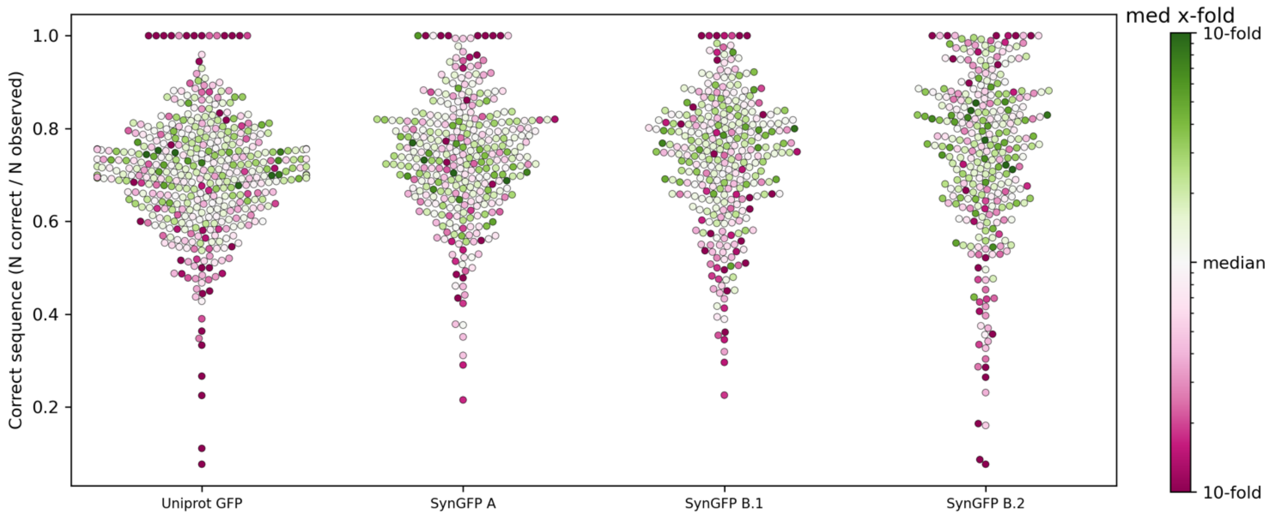


**Supplementary Figure 9. Fraction of correct assemblies without synthesis error.** Due to oligo synthesis error, correct assemblies with all fragments assembled in the correct order may contain sequence errors. To assess the fraction of assemblies that are also sequence-correct, we used pbmm2 to align designed library sequences against PacBio reads. Sequences are plotted by the fraction of mapped reads that were perfect matches. Each sequence is colored by its abundance (x-fold from the median-abundant sequence) on a log scale. More than 60% of mapped reads are error-free for abundant sequences (>= median abundant sequence).

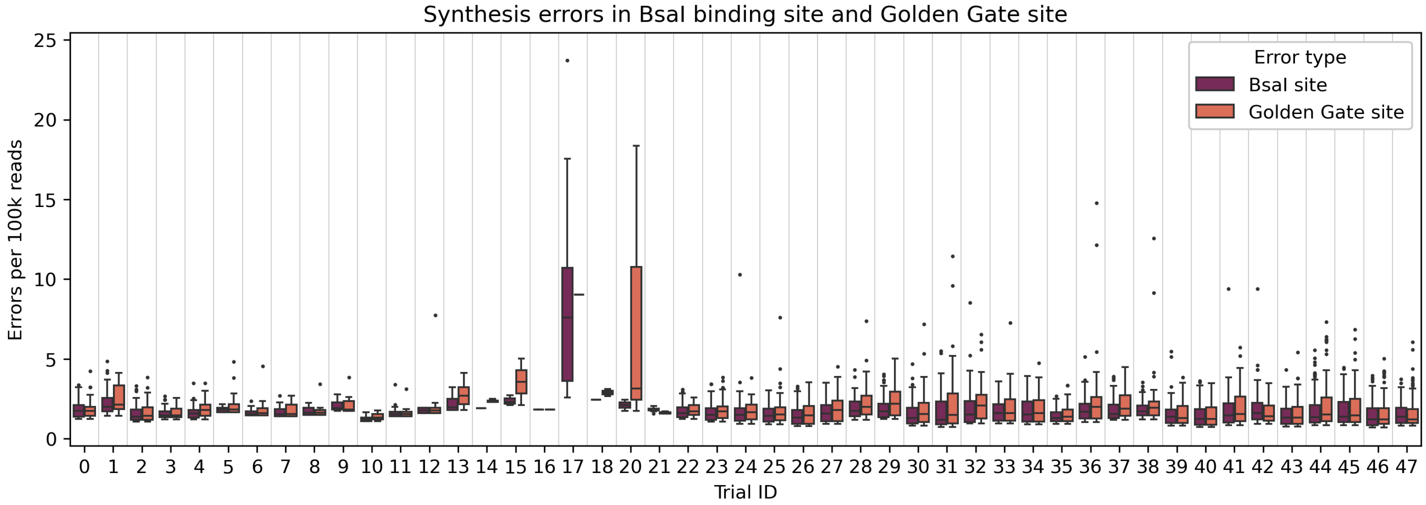


**Supplementary Figure 10. Synthesis errors in the BsaI binding site or Golden Gate site.** We applied Illumina sequencing for amplified oligos from parameterization assemblies. We report errors as the total number of reads normalized to 100k that contain synthesis errors in either the BsaI binding site or Golden Gate site. In most cases, synthesis errors in the BsaI and GG sites occur in < 0.005% of reads.

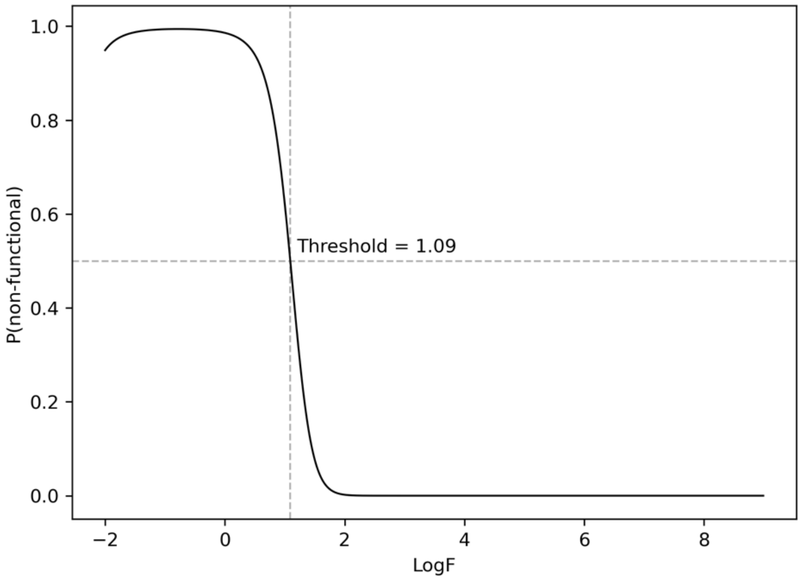


**Supplementary Figure 11. LogF functional cutoff for GFP fluorescence.** A Gaussian Mixture Model from the scikit-learn[2] package was fit to designed sequences with >= 55 cumulative counts across sorted populations (N=314). We used LogF values averaged across two experimental replicates. We plotted P(non-functional) for LogF values in increments of 0.01 from -2 to 9. The LogF closest to 0.5 was used as the functional cutoff.

**Table S1. Design statistics reporting number of observed and functional sequences in SynGFP.** Designed indicates the number of sequences included in the library. "> 55 counts" is the number of sequences observed >= 55 times across all sort bins. N functional are the number of sequences above the LogF cutoff 1.09. Fraction functional is the fraction of observed designs that are functional.

| Population | Designed | > 55 counts | N Functional | Fraction functional |
|---|---|---|---|---|
| Negative | - | 76 | 2 | .03 |
| MIFST-doubles | 15 | 15 | 12 | .80 |
| CARP-640M-doubles | 15 | 14 | 5 | .36 |
| MIFST-MCMC | 100 | 85 | 0 | 0 |
| CARP-MCMC | 100 | 90 | 0 | 0 |
| ESM-MSA-MCMC | 116 | 109 | 66 | .61 |
